# Advancing rust resistance in elite wheat with haplotype mapping and a novel introgression strategy

**DOI:** 10.1101/2025.09.22.677963

**Authors:** Seema Yadav, Shannon Dillon, Meredith McNeil, Eric Dinglasan, Dilani Jambuthenne, Rohit Mago, Peter N. Dodds, Lee T. Hickey, Ben J. Hayes

## Abstract

Wheat production is continually threatened by stripe and leaf rust because virulent races rapidly overcome single race-specific genes. Durable, broad-spectrum resistance is needed. Adult plant resistance (APR) provides partial, stable resistance from multiple minor-effect loci acting additively; pleiotropic loci like *Lr34/Yr18* and *Lr46/Yr29* add durability. We used an elite Australian panel (OzWheat=589) and a diverse landrace panel (Vavilov=295), genotyped with ∼30K SNPs and phenotyped across environments. Linkage disequilibrium partitioning defined 7,659 genome-wide haploblocks. To prioritise robust signals, we ranked haploblocks by haplotype effect variance and examined the top 100 per trait. For stripe rust, 52/100 were significant, with 32 shared across panels; for leaf rust, 50 were significant, 29 also detected in Vavilov. Several intervals co-localised with APR regions (*Lr46*/*Yr29*), and one 7BL interval intersected seedling gene *Lr14a*. To translate mapping into breeding decisions, we developed an introgression fitness index to quantify the value of resistant haplotypes in elite backgrounds. Using elite cultivar Scepter, we applied a genetic algorithm to select 50 donor parents carrying desirable haplotypes. Simulations showed that pyramiding these haplotypes can enhance resistance while maintaining elite genomic background. This study provides practical breeding tools, including haplotype catalogue and a novel selection index to accelerate rust-resistant wheat development.

## Introduction

Wheat (*Triticum aestivum* L.) is a major global staple, contributing nearly 20% of the world’s caloric intake (Shiferaw et al. 2013). Its production is persistently threatened by fungal rust diseases, notably stripe rust (*Puccinia striiformis* f. sp. *tritici*) and leaf rust (*P. triticina*), which are among the most damaging pathogens worldwide (Figueroa et al. 2018; Savary et al. 2019). Stripe rust can reduce yields by up to 60% in susceptible varieties under conducive conditions (Ma et al. 2025; Van-Zivkovic et al. 2025), whereas leaf rust typically causes lower (e.g. 1-20%) but still economically significant losses (Figueroa et al. 2018; Huerta-Espino et al. 2011). Stripe rust is favoured by cooler, moist conditions, while leaf rust generally prefers moderate to warm temperatures. Disease pressure and geographic range have expanded in recent decades, influenced by climate warming and increased movement of inoculum through trade and travel (Figueroa et al. 2018).

Rust pathogens exhibit rapid evolutionary dynamics, with frequent emergence of new virulent races that overcome deployed resistance genes, highlighting the urgent need for broad-spectrum and durable resistance in wheat (Chen and Kang 2017; Singh et al. 2015). Breeding strategies combine all stages (seedling) resistance, typically race-specific and conferred by single major genes, with adult plant resistance (APR). APR is expressed later in development, is usually partial and broad-spectrum, and tends to be more durable across diverse field conditions. However, APR is quantitatively inherited, environmentally influenced, and often involves many minor effect loci, complicating gene discovery and deployment (Van-Zivkovic et al. 2025; Wang et al. 2023). Despite these challenges, several APR genes, such as *Lr34/Yr18* and *Lr46/Yr29*, have been widely used to enhance durable rust resistance (Ellis et al. 2014; McIntosh et al. 2018). These loci each provide only partial protection, but when pyramided with other genes, they can greatly improve the durability and breadth of resistance. Modern breeding programs, therefore, prioritise combining APR genes to achieve robust multi-rust resistance (Wang et al. 2023).

Genome-wide association studies (GWAS) using single-nucleotide polymorphisms (SNPs) are standard for identifying APR loci. However, SNP-based GWAS has limitations: low detection power for small-effect loci, testing thousands of SNPs incurs a high multiple-testing burden, and inability to capture multi-allelic variation (Bhat et al. 2021). These issues are pronounced in wheat, a polyploid species characterised by extensive linkage disequilibrium (LD) and genomic redundancy. Haplotype-based association mapping improves detection by modelling linked SNPs as multi-allelic haplotype blocks. This approach increases statistical power and reduces the number of tests compared to single-SNP analyses. Haplotypes enhance mapping resolution by delineating inherited genomic segments more precisely, capturing rare or complex variants missed by single SNPs (Lisker et al. 2022; Liu et al. 2020). However, the extent of improvement remains constrained by the underlying LD patterns and the size of haplotype blocks. In wheat, haplotype-led analyses have already demonstrated superior performance in identifying QTLs for disease resistance and other complex traits (Brinton et al. 2020; Cheng et al. 2024; Liu et al. 2020; Tong et al. 2024; Van-Zivkovic et al. 2025). Importantly, these methods are particularly well-suited to capturing functional variation within the extended LD blocks and homoeologous regions typical of polyploid genomes (Voss-Fels et al. 2019).

A rich but underutilised source of rust resistance alleles is exotic germplasm such as landraces (Jambuthenne et al., 2022; Riaz et al., 2018). Recent haplotype-based studies in historical wheat collections (e.g., Watkins and Vavilov panels) have uncovered unique resistance haplotypes that are absent in modern cultivars. Introgressing these haplotypes can improve resistance and agronomic performance in elite backgrounds (Cheng et al. 2024; Tong et al. 2024). These findings reinforce the potential of haplotype-led approaches to unlock novel allelic variation for breeding. Beyond diversity panels, haplotype-based methods have shown promise in breeding populations as well, enabling multi-environmental QTL detection for stripe rust resistance (Van-Zivkovic et al. 2025) and improving genomic predictions for leaf rust in hybrid wheat (Liu et al. 2020). Together, this highlights the expanding role of haplotype mapping in both pre-breeding and advanced cultivar development.

However, most studies to date focus on either exotic or elite germplasm, limiting direct breeding application. Resistance haplotypes identified in exotic germplasm are rarely compared to or transferable into elite breeding material. Thus, we often do not know whether a “novel” QTL found in landraces is truly absent from modern germplasm or simply undetected therein; conversely, some valuable alleles in landraces may already exist (or have analogues) in cultivars. Bridging this translational gap requires a unified genotypic and phenotypic framework spanning both exotic and advanced breeding lines. As noted in the Watkins study (Cheng et al. 2024), leveraging landrace diversity for crop improvement demands appropriate resources and informatics to bridge the gap between landrace alleles and modern breeding.

In this study, we address this gap by constructing a high-density haplotype map from a combined panel of Australian cultivars and breeding lines (OzWheat) and landraces from the N.I. Vavilov Institute. While association mapping primarily focuses on the OzWheat panel, a representative core set of 50 Vavilov lines was phenotyped in parallel to validate haplotype effects and identify novel resistance alleles absent in elites. By integrating exotic and elite germplasm in one framework, this study aims to: (i) identify genomic regions associated with stripe and leaf rust resistance using high-resolution haplotype mapping in the elite panel, augmented by landrace validation; (ii) compare the distribution and effect sizes of resistance-associated haplotypes between elite and exotic lines; (iii) develop a haplotype-introgression fitness index to prioritise donor parents for breeding; (iv) use simulations to evaluate strategies for stacking haplotypes and improving resistance.

This work offers new insights into the genetic architecture of rust resistance and provides practical resources to enhance durability in future wheat varieties. We demonstrate a translational pipeline from haplotype discovery to marker-informed introgression, which can serve as a model for other complex traits and crop systems.

## Material and methods

### Plant materials and study design

This study evaluated APR to stripe and leaf rust using two wheat panels. First, the Vavilov panel, comprised 295 landrace accessions representing global diversity (Tong et al. 2024). Second, the OzWheat panel, included 589 advanced breeding lines, landraces and modern cultivars representative of Australian germplasm (Hyles et al. 2024). For genome-wide association mapping, the OzWheat panel served as the primary population due to its larger sample size and comprehensive phenotypic dataset. In parallel, a Vavilov core set (n=50) was selected for phenotyping to validate haplotype effects and to identify resistance haplotypes absent from elite material. The core set was chosen to maximise allelic diversity and ensure broad genetic representation of the full Vavilov panel (Supplementary Fig. S1).

### Genotyping, data integration and imputation

All 295 Vavilov lines were genotyped using the Illumina iSelect 90K SNP array (Wang et al. 2014) and DArTseq (Kilian et al. 2012), resulting in ∼35,000 high-quality polymorphic markers (Tong et al. 2024). The OzWheat panel was genotyped using multiple SNP platforms, which we managed as two sub-cohorts: (i) OzWheat V1 (n = 299) had a composite high-density SNP dataset (∼115,260 SNPs) aggregated from the 90K array (∼21,559 SNPs), the Axiom 40K Breeder’s array (∼10,417) (Allen et al. 2017), DArTseq (∼20,062), and transcriptome-derived SNPs (∼63,222). (ii) OzWheat V2 (n = 289) was genotyped exclusively with the Illumina 90K SNP array (∼21,500 markers).

To ensure consistency across genotyping platforms and minimise imputation artefacts, genotype data from each panel were processed independently and aligned to the wheat reference genome (IWGSC RefSeq v1.0). We adopted a tiered imputation strategy to unify the datasets while balancing cost and resolution. After quality control, removing monomorphic and duplicate sites, markers with >20% missing data, and variants with minor allele frequency (MAF) ≤□5%, the high-density OzWheat V1 set (∼97k SNPs) was phased and used as an internal reference to impute missing or untyped markers in OzWheat V2 and the Vavilov panel. Before imputation, marker identities were matched by physical position; strand and allele nomenclature (reference/alternate) were standardised; and C/G ambiguous sites were checked for strand flips. Samples with >50% missing genotypes were also excluded. Filtering and indexing of VCF files were conducted using bcftools (bcftools/1.15.1-gcc-11.3.0) and vcftools. To assess allele frequency concordance between reference and target panels, we compared MAFs for overlapping SNPs and removed markers with large frequency discrepancies (Supplementary Fig. S2 and S3). This filtering step retained 14,267 SNPs for OzWheat V2 and 12,137 SNPs for the Vavilov panel.

Genotype imputation and haplotype phasing were performed chromosome-by-chromosome using Beagle v5.4 (build 18 Mar 2022), which simultaneously infers haplotype structure and fills in missing genotypes through localised LD-based clustering (Browning et al. 2021). The imputation quality was evaluated using Beagle’s dosage R^2^ metric (DR^2^, which reflects the squared correlation between imputed and true allele dosage), and only high-confidence markers (DR^2^ ≥ 0.8, a stringent threshold chosen to minimise imputation error) were retained, yielding ∼49,000 SNPs for OzWheat V2 and ∼42,000 SNPs for the Vavilov panel. Following imputation, the datasets were joined to construct a unified genotype matrix. Prior to merging, SNP coordinates (chromosome and base-pair position) were verified and ordered by chromosome and position. Allele designations (REF/ALT) were standardised across panels, and any metadata discrepancies were resolved. The final merged set comprised 29,972 high-confidence SNPs shared across n□=□866 lines (OzWheat□+□Vavilov), providing the foundation for downstream haplotype analyses and association testing.

### Field trials and disease assessment

Field trials were conducted over two growing seasons (2023 and 2024) across five locations in major Australian wheat-growing regions to capture diverse environments and different pathotypes. Stripe rust nurseries were at DAF Hermitage (Warwick, QLD), AgVic Horsham (VIC) and PBI Cobbitty (NSW) in both years. Leaf rust nurseries were established at DPIRD Shenton Park (WA) and PBI Cobbitty (NSW) in 2023, with AgVic Horsham added in 2024 to extend environmental coverage. In total, phenotypes were obtained from 584 OzWheat lines (5 excluded due to seed unavailability) and the 50-line Vavilov core set. Trials used a randomised complete block design with two replicates per site. Local agronomic practices and spreader rows were used to ensure uniform and high disease pressure, with natural infections supplemented by inoculation where necessary (Supplementary Data).

Adult plant rust response was assessed on a standardised 1-9 ordinal scale (1= highly resistant [R], 2 = RMR, 3 = MR, 4 = MRMS, 5 = MS, 6 = MSS, 7 = S, 8 = SVS, 9 = VS very susceptible [S]). At the Western Australian leaf rust trials (DPIRD, Shenton Park), disease notes were initially recorded as a percentage coefficient of infection scale (0-100) following local protocols. These percentage scores were converted to the 1-9 scale for comparison across locations, using the method described by Bariana et al. (2007). Rust scores were evaluated on a weekly basis starting from the emergence of the flag leaf and continued until the susceptible control reached a disease score of 8 to 9. For analysis, the final disease rating recorded for each line was used. Raw phenotype data were collected collaboratively and stored in the CSIRO Data Access Portal (DAP, https://data.csiro.au/collection/csiro:61604).

**Table 1.**
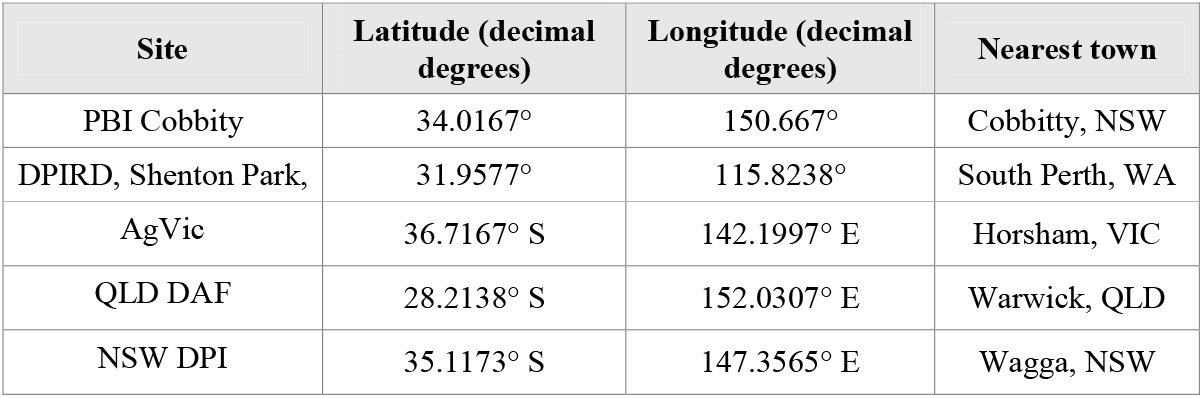
Locations of field trial sites used for rust resistance phenotyping.

### Phenotypic data processing and statistical analysis

Prior to statistical analysis, phenotypic data were subjected to quality control to assess missing data, outliers, and consistency across locations and years. For each trial, a linear mixed model was fitted to obtain best linear unbiased estimates (BLUEs) for downstream genetics. For each trial, genotype and replicate were treated as fixed effects in the following model:

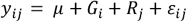

where *y*_*ij*_ is the observed disease score for genotype *i* in replicate *j, μ* is the intercept, *G*_*i*_ is the fixed effect of genotype *R*_*j*_ is the fixed effect of replicate, and *ε*_*ij*_ is the residual error term. To evaluate the reliability of each trial, genotype was treated as a random effect to capture genetic variance and calculate broad-sense heritability (*H*^*2*^) on a line-mean basis. Heritability was calculated as:

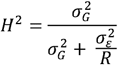

where 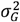 is the genotypic variance, 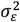 is the residual variance, and R is the number of replicates.

A single-stage multi-environment trial analysis was conducted to estimate genotype performance across trials. Although disease scores were recorded on an ordinal 1-9 scale, they were treated as approximately continuous for analysis, following standard practice in field pathology. Combined-site-year BLUEs were computed for each genotype by fitting a linear mixed model across all trials, with genotype as fixed and trial (defined as a location-year combination) as a random effect with heterogeneous residual variance by trial using the following model.

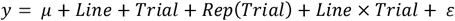

The resulting genotype BLUEs across environments were used for downstream haplotype analyses.

### Population structure and linkage disequilibrium analysis

Population structure was assessed using a unified genotype matrix (29,972 SNPs; n = 866 lines) described above. Principal component analysis (PCA) was performed on a genetic distance matrix derived from the combined OzWheat + Vavilov dataset using the *SelectionTools* R package (v21.3; population-genetics.uni-giessen.de/∼software/). and base R, with genotypes coded as minor allele dosages (0/1/2), mean-centred per marker and scaled to unit variance. To summarise group structure, k-means clustering (K = 5) was applied to the PCA coordinates.

LD decay was quantified within each chromosome. Pairwise LD was calculated as the squared allele frequency-correlation (*r*^*2*^) between adjacent SNP pairs from the imputed genotype dosages. For each chromosome, *r*^*2*^ values were plotted against physical distance (Mbp), and a LOESS curve was fitted in R to describe average decay. A genome-wide LD decay curve was generated by pooling chromosome-wise LD estimates. LD extent was summarised by reporting the approximate physical distance at which the smoothed *r*^*2*^ declined to reference thresholds (e.g., *r*^*2*^ = 0.2). The R package ggplot2 (Wickham 2016) was used to visualise the results, including a locally estimated scatterplot smoothing (LOESS) line.

### Haplotype block construction and QTL mapping using a local GEBV approach

#### LD-based haplotype block construction

Genome-wide haplotype blocks (haploblocks) were defined based on LD using the *SelectionTools* R package. The process was performed separately for each of the 21 wheat chromosomes to account for differences in recombination and population structure. Starting from the leftmost SNP of a chromosome, adjacent markers were sequentially grouped into a block if their pairwise LD (measured as *r*^*2*^) between the new SNP and the current block’s boundary (the last SNP in the block) exceeded a set threshold. We chose an *r*^*2*^ threshold of 0.5, adopted from Tong et al. (2024). The tolerance parameter was set to t = 0, meaning no low-LD exceptions were allowed; every SNP added to a block had to show *r*^*2*^ *≥* 0.5 with the block’s flanking SNP. This stringent criterion ensures that all SNPs within a block are in strong LD with each other. If this condition was not met, the block was finalised, and a new block was initiated. Markers not in LD with any neighbouring SNP (*r*^*2*^ < 0.5 on both sides) were treated as individual haploblocks. For each haploblock, we recorded the physical start and end positions, number of included SNPs, and block size for downstream analyses.

#### Panel-specific analysis strategy

To ensure consistent haplotype definitions across datasets, haploblocks were defined using the combined genotypic data from all 866 lines. However, because phenotypic data were only available for 589 OzWheat panel and a 50-line Vavilov core set, all downstream analysis, including SNP effect estimation, haplotype effect calculation, and haploblock discovery, were conducted separately for these two phenotyped subsets.

#### Estimation of SNP effects and block-wise local genomic breeding values

To estimate SNP effects on the rust resistance traits, we applied a ridge regression best linear unbiased prediction (RR-BLUP) model (Endelman 2011), implemented in the *SelectionTools* R package. This whole-genome regression approach includes all SNP markers simultaneously as random effects. The model is defined as:

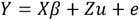

Where, *Y* is the vector of phenotypic values (e.g. across-environment BLUEs for stripe rust or leaf rust), *X* is the design matrix for fixed effects (intercept), *β* is the vector of fixed effects, *Z* is the design matrix allocating observations to SNP effects, *u* is the vector of random SNP effects with 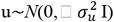 and *e* is the residual error vector following 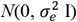. Variance components and the equivalent ridge penalty 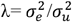 were estimated by REML. RR-BLUP shrinks individual SNP effects towards zero, which is appropriate given that we expect many small-effect loci. To derive haplotype effects, we computed the additive contribution of SNPs within each LD-defined haploblock (Tong et al. 2024; van den Berg et al. 2019; Voss-Fels et al. 2019).

For each haploblock *b*, the local genomic estimated breeding value (local GEBV) for line *i* was calculated as: 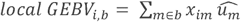

where *x*_*im*_ is the allele dosage for line *i* at marker *m*, and 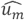 is the estimated effect of marker m. These block-wise breeding values capture the cumulative additive effect of each haplotype on the trait of interest (Kemper et al. 2012; Varshney et al. 2022; Xiang et al. 2021).

Following haplotype effect estimation, a haplotype catalogue was developed for the OzWheat panel, identifying which haplotype allele was associated with resistance (low disease score) and which lines possessed that allele.

#### Identification of candidate QTL blocks

To identify genomic regions potentially associated with rust resistance traits, the variance of haplotype effects within each LD-defined block was used as a heuristic indicator of QTL. The underlying rationale is that if a haploblock harbours a QTL, the haplotypes within that block will exhibit divergent effects: some conferring resistance, others neutral or susceptible, resulting in elevated variance in local GEBVs (van den Berg et al. 2019; Xiang et al. 2021). Conversely, haploblocks with no true effect on the trait should show only small random differences in haplotype effects (low variance). Following the approach of Kemper et al. (2012), a haploblock was declared as a genomic region associated with rust traits if the variance among its haplotype effects was at least 100 times greater than the genome-wide average haplotype effect variance. This threshold is deliberately stringent and highlights only those regions where haplotype effects are much more dispersed than the genome-wide background and serves for prioritisation. Additionally, haploblocks were ranked by their haplotype effect variance, and the top 100 blocks per trait were examined. To statistically assess the contribution of the top 100 haploblocks to trait variation, a chi-square test was performed. The null hypothesis assumed that haplotype effects within a block explain zero genetic variance. Under this assumption, the ratio of haplotype effect variance to total genetic variance (from the RR-BLUP) should approach zero. Specifically, for a given block, the test statistic was:

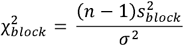

Where 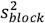 is the variance of haplotype effects within the block (local GEBV variance), *σ*^2^ is the total genetic variance for a particular trait estimated from the RR-BLUP model, and *n* −1 is the degree of freedom (with *n* being the number of haplotypes observed at that block). The *p*-value for each block was calculated as: 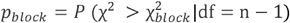. Blocks with significant *p-*values (after appropriate multiple-testing correction) were considered significant genomic regions influencing rust resistance.

#### Haplotype diversity in OzWheat and Vavilov core set

For each haploblock *b* and panel *p* (OzWheat, Vavilov core set), the Shannon diversity index was computed as:

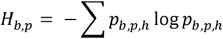

Where *p*_*b,p,h*_ is the frequency of haplotype h in panel p. Diversity was summarised per panel across blocks. Panel differences were tested via the Wilcoxon rank-sum on block-wise H values.

#### Partitioning genetic variance by haploblock

The contribution of each LD-defined haploblock to rust resistance was quantified using the GVCHAP software (Prakapenka et al. 2020). First, we partitioned additive genetic variance between genome-wide SNPs and LD-defined haplotype blocks using GVCHAP, fitting the model (SNP + haplotype additive values):

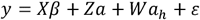

where y is the vector of across-environment genotype BLUEs for stripe rust, *Z and W* are incidence matrices for SNP genotypes and haplotypes states, 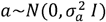, and 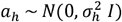 and 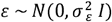 SNP effects, haplotype effects and residual. Variance components 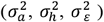 were estimated by REML. GVCHAP outputs per-marker and per-haploblock “heritability” metrics, defined as the proportion of phenotypic variance explained by the corresponding random effect conditional on the background terms. For a haploblock *b*, the block-level contribution is 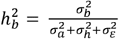, where 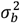 is the variance attributed to block *b* within 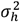 . We interpret 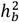 as how much genetic variation is captured by that block, given the genome-wide SNP background and the other haplotype terms included in the model. The overall contribution of haplotypes vs. SNPs to heritability by evaluating three models: (i) SNPs only, (ii) haplotypes only, and (iii) a combined SNP + haplotype model.

### Parent selection via a haplotype-based introgression fitness index

To guide parent selection for an introgression program, a custom fitness function was developed that ranks lines based on their complement of favourable haplotypes relative to a benchmark elite cultivar. In our case, ‘Scepter’ (widely grown in Australia, known for good yield and moderate rust resistance) was chosen as the reference genotype. For a candidate set of *n* parents *S* (selected set = 50 parents as a group) and for a given trait *t*, the fitness function is defined as:

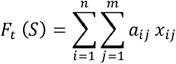

For donor line *i*, define the indicator:

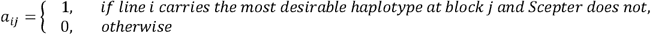

*x*_*ij*_ is the estimated effect of the haplotype carried by parent *i* at block *j, n* is the number of candidate parents (| *S* | = 50), *m* is the number of haplotype blocks considered, in this case, the entire genome-wide set (7,659 blocks).

The scoring function prioritises lines that contribute novel, favourable alleles absent from Scepter, rather than simply selecting the most resistant individuals. The procedure was applied independently to each trait, and final selection was done by assigning equal weights (50% Yr, 50% LR) to both traits. Parent selection was formulated as a constrained combinatorial optimisation problem and solved using a genetic algorithm to maximise the fitness function (Yadav et al. 2025). To ensure genetic diversity, selected lines were evaluated using PCA on genome-wide markers. Additionally, GEBVs for both rust traits were visualised to confirm that selected lines included not only highly resistant genotypes but also moderately or less resistant ones that carry unique favourable haplotypes absent in Scepter.

#### Crossing design and simulation for progeny performance

To evaluate the potential genetic gain in rust resistance through haplotype introgression, we designed a crossing-simulation framework as part of a pre-breeding strategy we term *RustHapSelect*. The goal was to assemble a pre-breeding pool of individuals that stacked as many favourable resistance haplotypes as possible from selected donor lines together with elite cultivar background haplotypes from ‘Scepter’. From the 50 selected donor lines, each was ‘crossed’ with Scepter *in silico* to generate 50 single-cross F □ families. To stack haplotypes from multiple donors into a new pool pre-breeding material, we then simulated all pairwise intercrosses among the 50 F □s, resulting in 1,225 double-cross (DC_1_F_1_) combinations. From this base, we implemented two contrasting and commonly used crossing strategies:

##### Scenario 1: Forward Breeding (Selfing Strategy)

For each of the 1,225 DC_1_F_1_ combinations, 50 progenies were simulated (totalling 61,250 individuals). These progenies underwent three generations of selfing (DC_1_F □ → DC_1_F □ → DC_1_F □). At DC_1_F_4_, each family was scored as the mean of (i) the family average index and (ii) the mean of the top 5 individuals. Families were ranked by this score; the top 50 DC families were selected for pre-breeding crossing activities.

##### Scenario 2: Backcrossing to Elite Parent

Starting from the same 1,225 DC_1_F_1_ combinations, 50 progeny per family were generated, as in Scenario 1. These were backcrossed once to Scepter, to produce 10 BC □F_1_ progeny per DC family. The BC □F_1_ progenies were advanced by three generations of selfing from BC □F □ → BC □F □. DC families were ranked using the same selection index as Scenario 1. Similarly, the top 50 DC families were selected to guide subsequent breeding efforts.

Although the crossing design aimed to combine favourable resistance haplotypes with elite background haplotypes from Scepter, for simplicity, the simulations were performed using genome-wide SNP data. As such, the number of retained resistance or Scepter background haplotypes was not explicitly tracked. Progeny performance was evaluated based on the composite rust resistance index, which integrates leaf and stripe rust genetic values equally (50:50 weight). All simulations were run using the *AlphaSimR* R package (Gaynor, Gorjanc,Hickey 2021), which models recombination, segregation, and inheritance using user-defined genetic maps and founder haplotypes. Each simulation was run for 10 replicates to account for stochasticity in recombination and segregation. The top 50 DC families per scenario represent the most promising combinations for rapidly stacking resistance haplotypes into elite germplasm unere the *RustHapSelect* pre-breeding strategy.

## Results

### Phenotypic variation in rust resistance across diverse wheat germplasm

We evaluated stripe and leaf rust resistance using two wheat panels: OzWheat, representing elite Australian breeding lines, and Vavilov, a global landrace accession. These panels capture both modern adaptation and untapped diversity to identify novel resistance sources and genome-wide haplotype effects. Field phenotyping across environments revealed substantial variation in rust disease. Overall, data quality was high, especially for stripe rust, with < 1% missing data across most sites and years, indicating reliable disease pressure and scoring (Supplementary Fig. S4). For leaf rust, however, data completeness varied by site; notably, the PBI Cobbitty (NSW) trial had high missing data (22.3% in 2023; 36.3% in 2024), and the Horsham (AgVic) 2024 trial had ∼30% missing values (Supplementary Fig. S4).

Rust scores on the 1-9 scale spanned the full range in most cases, reflecting broad genetic variation. Stripe rust severity scores exhibited a continuous distribution from high resistant ([R-RMR] ∼ 1-2) to highly susceptible (∼8-9) across locations (Fig. 1). Leaf rust scores, however, showed greater variability between sites and were often skewed toward resistance (Fig. 1), particularly at PBI, likely due to confounding from co-occurring stripe rust infections that hindered accurate leaf rust scoring. At Shenton Park (WA), leaf rust ratings ranged across the scale, suggesting clear separation of genotypes under local epidemic conditions.

**Fig. 1:**
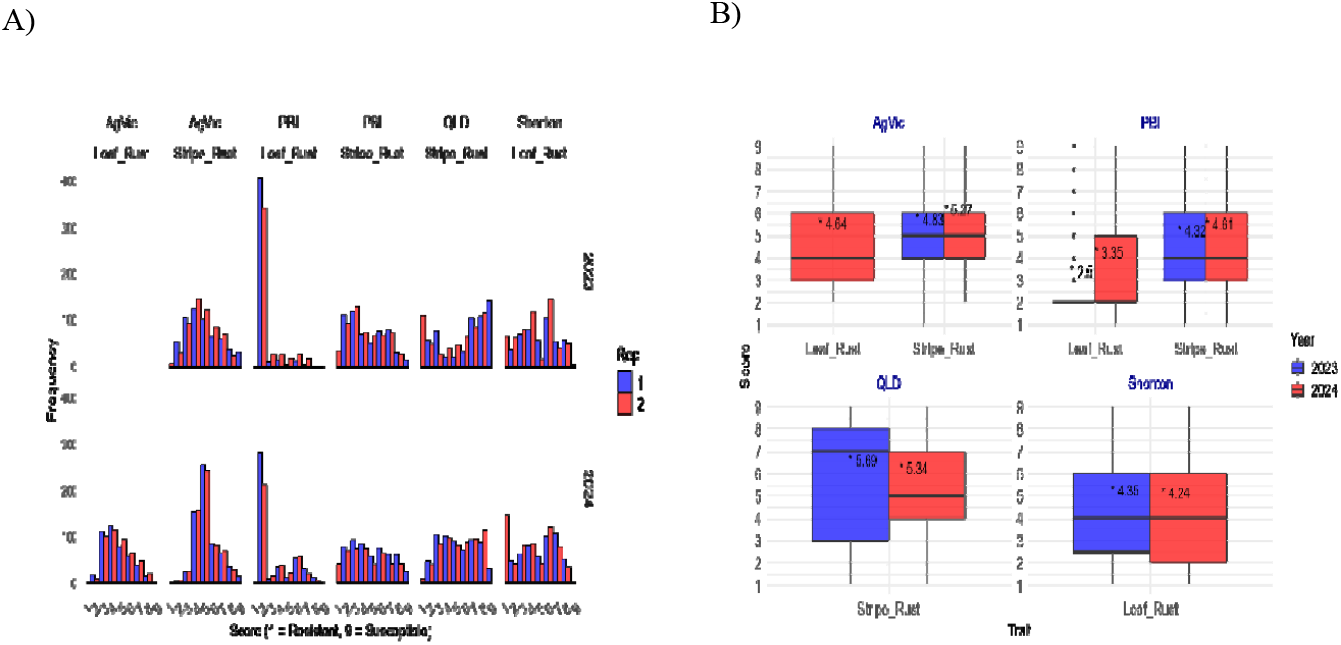
**A)** Frequency distribution of leaf rust and stripe rust scores (1-9 scale; 1= resistant, 9 = susceptible) for the combined germplasm, across field trial locations (AgVic, PBI, QLD, and Shenton) for 2023 and 2024. Stripe rust shows a wide range of severity in all considered locations. For leaf rust, the Shenton Park (WA) data exhibit good spread, while at PBI (NSW), most of the lines are scored as resistant. **B)** Boxplots of rust scores by location and year.

Descriptive statistics summarising disease severity are presented in Supplementary Table S1. Mean stripe rust scores ranged from 4.32 (MR-MRMS) at PBI Cobbitty to 5.69 (MS) at Hermitage (QLD) in 2023, with standard deviations reflecting site-specific disease intensity (e.g. 2.78 at QLD) (Fig. 1B). For leaf rust, mean scores ranged from 2.50 (RMR) at PBI in 2023 to 4.64 (MR-MRMS) at AgVic in 2024 (Fig. 1B). Standard deviations for leaf rust were generally higher at Shenton and PBI, indicating broader phenotypic variation (e.g. 2.34 at Shenton in 2024) (Supplementary Table S1).

Broad-sense heritability (H^2^) for stripe rust was consistently high (H^2^ > 0.85), supporting the reliability of phenotypic data (Supplementary Fig. S5). This lends confidence that the stripe rust BLUEs accurately capture genetic differences, providing a solid basis for association mapping. Leaf rust heritability was more variable; moderate at PBI (H^2^ ∼ 0.5), likely due to scoring noise from stripe rust co-infection, and higher at Shenton Park (Supplementary Fig. S5), indicating greater reliability. Given these observations, we proceeded with a conservative strategy for leaf rust analysis, focusing on the most reliable environment (Shenton Park) for primary QTL discovery and using other sites as supporting information only.

Multi-environment modelling indicated significant genotype effects for both rusts, with notable Line × Trial (Location-Year) interactions. Location explained more variation than year, consistent with differences in local epidemics and/or pathotypes. Pairwise correlations of site-specific BLUEs for stripe rust were higher in 2023 (*r* = 0.57-0.83), indicating consistent genotype rankings across locations. In 2024, these correlations declined (*r* = 0.37-0.61), with weaker agreement involving PBI Cobbitty, suggesting that the stripe rust expression at PBI diverged somewhat, potentially due to microclimatic or race differences causing interactions.

### Genetic data integration, population structure, and LD patterns

After quality control and imputation, we obtained 29,972 high-confidence SNPs across 866 wheat accessions (OzWheat and Vavilov; see Methods). SNPs were distributed across all 21 chromosomes, though not uniformly. Subgenome B exhibited the highest marker density (14,052 markers), followed by subgenome A (12,463) and D (3,457), reflecting the known bias of array-based SNPs toward A/B genomes (Supplementary Table S2). Within subgenome B, the 90K SNPs platform contributes the most with 5,905 markers, followed by Transsnps with 5,637 markers. In contrast, subgenome D has the fewest markers, with a substantial proportion, 1,591, derived from Transsnps. Genome-wide coverage was visualised as SNP counts per 1-Mb window (Fig. 2), revealing densities ranged from 0 to >136 SNPs per Mb. The highest coverage was observed on chromosomes 1A, 5B, and 7B, indicating regions of particularly dense marker representation.

**Fig. 2:**
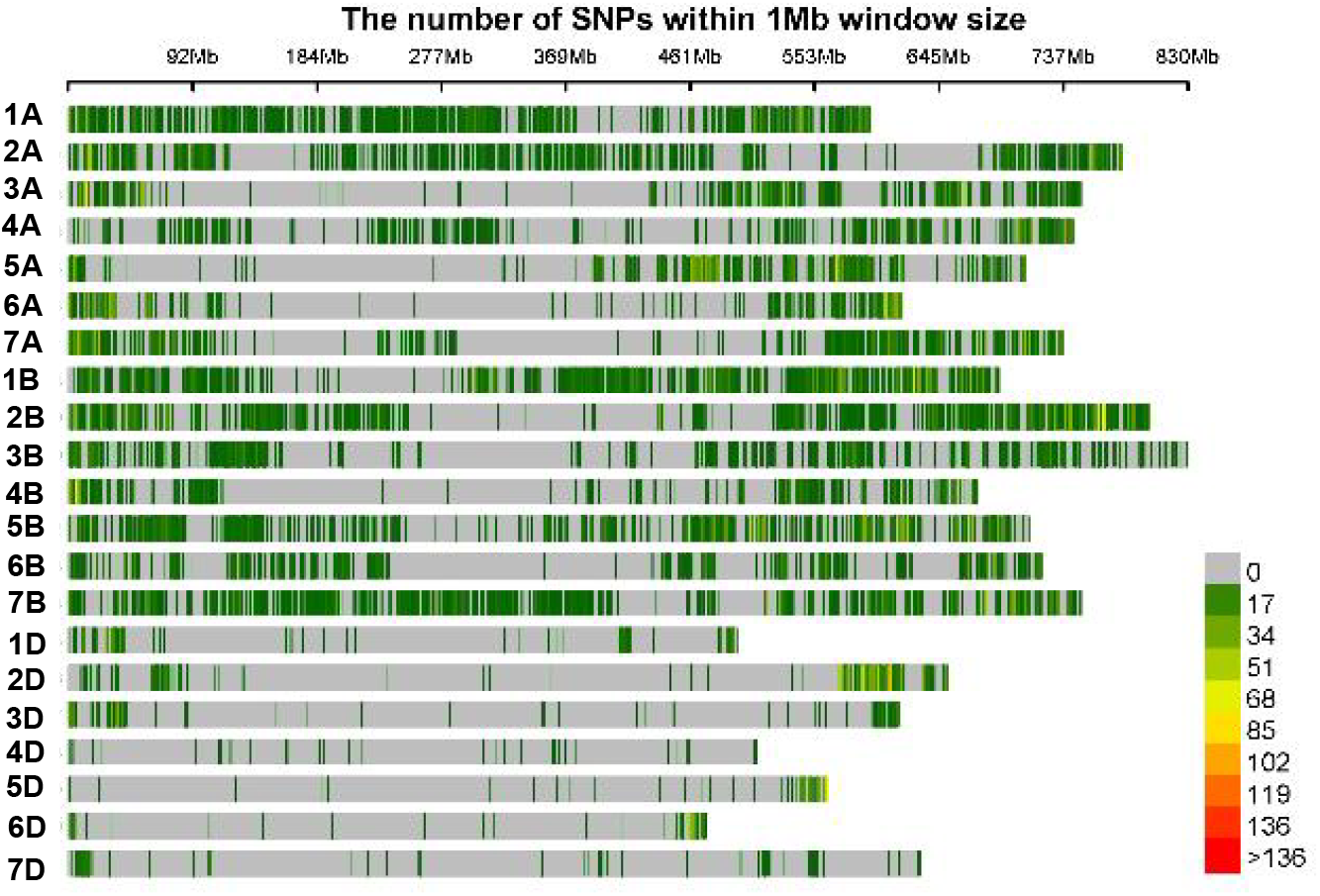
SNP density distribution across the wheat genome for the final integrated marker set (29,972 SNPs). SNPs are distributed in 1Mb intervals, colour-coded from light grey to red to indicate increasing densities.

Principal component analysis (PCA) revealed clear population structure among the combined panel (Fig. 3A). The first three PCs explained 15.14%, 10.04%, and 6.44% of genetic variance, respectively (Fig. 3A). Five major clusters were identified, with cluster1 predominantly representing Vavilov (108 +29 Oz) and cluster2 (n=103) exclusively OzWheat lines. Clusters 3-4 were composed mostly of OzWheat lines, but with additional sub-structure likely related to the breeding program or pedigree.

**Fig. 3:**
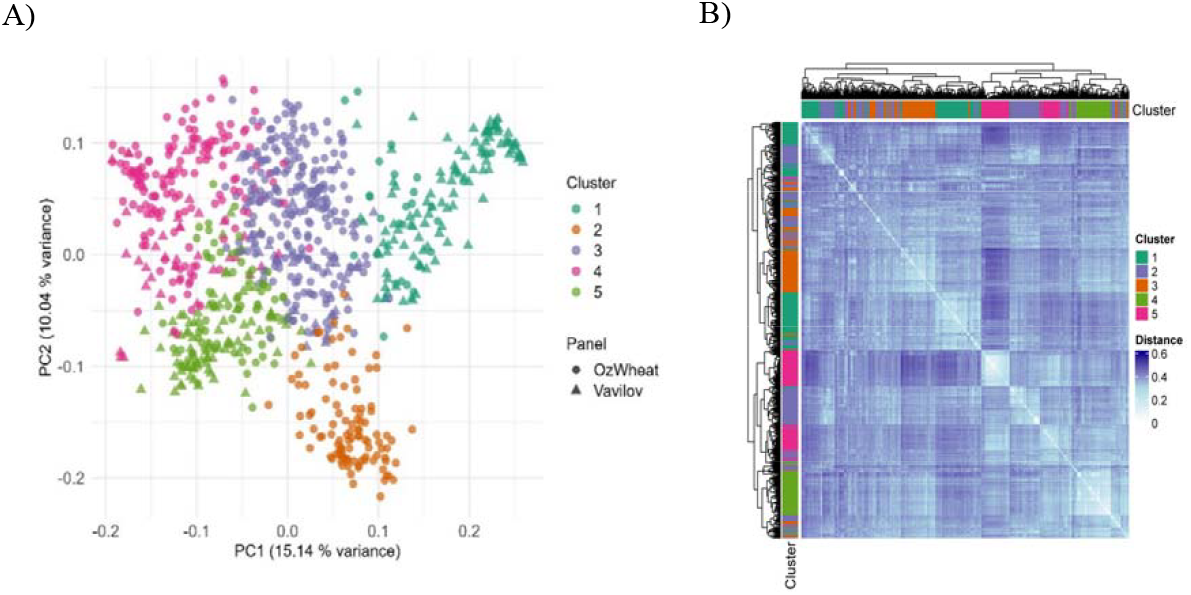
Population structure in the combined population. **A)** PCA analysis of combined wheat accessions (∼866 lines) using genome-wide SNPs. Samples are grouped into five clusters and represent two genetic panels: OzWheat (circles) and Vavilov (triangles). **B)** Heatmap of pairwise genetic distances among wheat accessions. Pairwise genetic distances are visualised as a heatmap, with lighter shades indicating greater similarity. Accessions are grouped into five clusters consistent with the PCA analysis. Rows and columns are ordered to highlight the substructure and relationship within and between clusters.

A heatmap based on pairwise genetic distance (Fig. 3B) revealed strong within-cluster genetic similarity, particularly for clusters 1 and 4, which formed compact, light-colored blocks. Greater genetic divergence (darker blue) was observed between clusters, notably between clusters 1 and 5. Some overlap between clusters 2 and 5 suggests potential shared ancestry. These patterns align with the PCA results and confirm clear population structure across the combined OzWheat and Vavilov panels.

LD decay was assessed using the unified marker set. In the OzWheat panel (=584), the LOESS-smoothed *r*^*2*^ curve declined to half its initial value (0.2) at ∼12 Mb (Supplementary Fig. S6), reflecting extended LD blocks likely due to breeding history and limited recombination. The Vavilov subset showed a comparable LD pattern, with slightly faster decay in some cases, reflecting its diverse origins. These LD profiles support haplotype block construction and association analysis, as many markers are inherited in large segments.

### Genomic regions associated with stripe and leaf rust resistance identified through haplotype analysis

#### Genome-wide haplotype-block discovery

Haplotype blocks were inferred from the integrated genotype data, applying an LD threshold *r*^*2*^ ≥0.5 with no tolerance for non-linked markers. In total, 7,659 blocks comprising 41,229 haplotypes were detected across the wheat genome. Block distribution varied across chromosomes, with 2B having the highest number of haplotype blocks (905), followed by 5B and 7A with 691 and 686 blocks, respectively. Chromosome 4D exhibited the lowest number of blocks (24), reflecting the sparse marker density (Fig. 4). On average, blocks contained ∼ 4 SNPs, but sizes varied widely. For example, 1D (known for lower diversity) had the largest average block size (14.12 SNPs per block), while 4D had the smallest (2.5 SNPs) (Fig. 4). This metric is not a proxy for physical size because marker density is uneven across the genome. Throughout, we therefore report block size by the physical coordinates of the flanking markers (Start-End position, Mb) as well as SNP count per block (Supplementary Data). Haploblock 7B:b006788 spanned 81.3 Mb with 275 SNPs, while 7D:b007213 had similarly high marker count (205 SNPs) in just 6.07 Mb, indicative of a region of high LD. Conversely, 6A:b002448 covered 85.8 Mb with only 11 SNPs, highlighting regional disparities in LD and marker coverage.

**Fig. 4:**
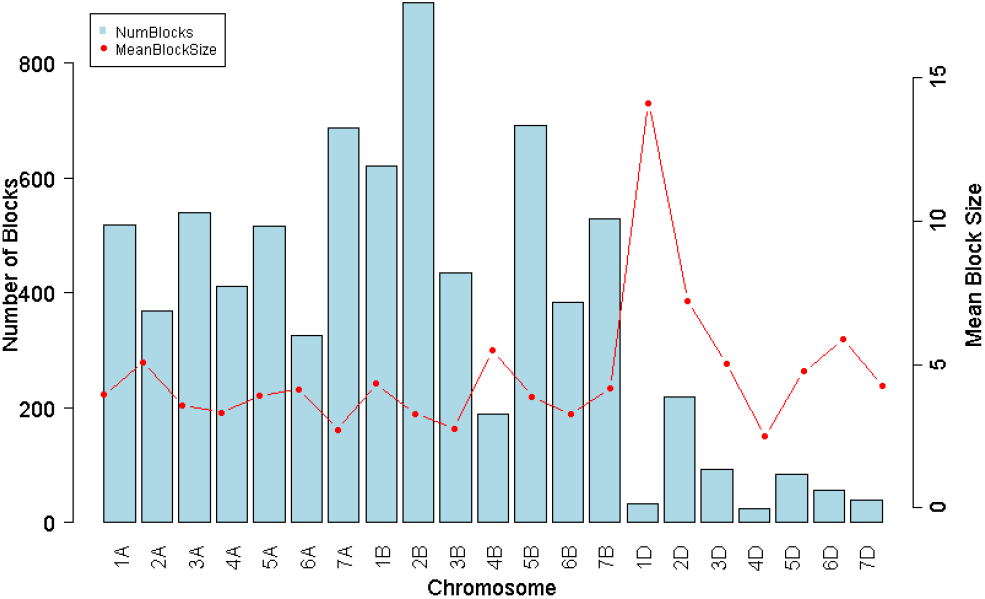
Total number of haplotype blocks (bars, left axis) and average number of SNPs per block (dots, right axis) across the genome.

#### Haplotype effects and identification of key resistance-associated regions

Haplotype blocks were ranked by haplotype-effect variance, and the top 100 per trait were tested for association (Fig. 5). For stripe rust, 52 blocks were statistically significant after chi-square testing with multiple testing correction. Of these, 32 blocks were identified in both OzWheat and Vavilov panels, despite separate association analyses, underscoring their robustness and overlap of key loci across both panels. Similarly, 50 blocks showed significant associations with leaf rust, and 29 of these were also detected in the Vavilov panel, highlighting a substantial overlap. Manhattan plots for leaf rust associations that met the stringent threshold are provided in supplementary Fig. (S8a & S8 b) for OzWheat and Vavilov, respectively.

**Fig. 5:**
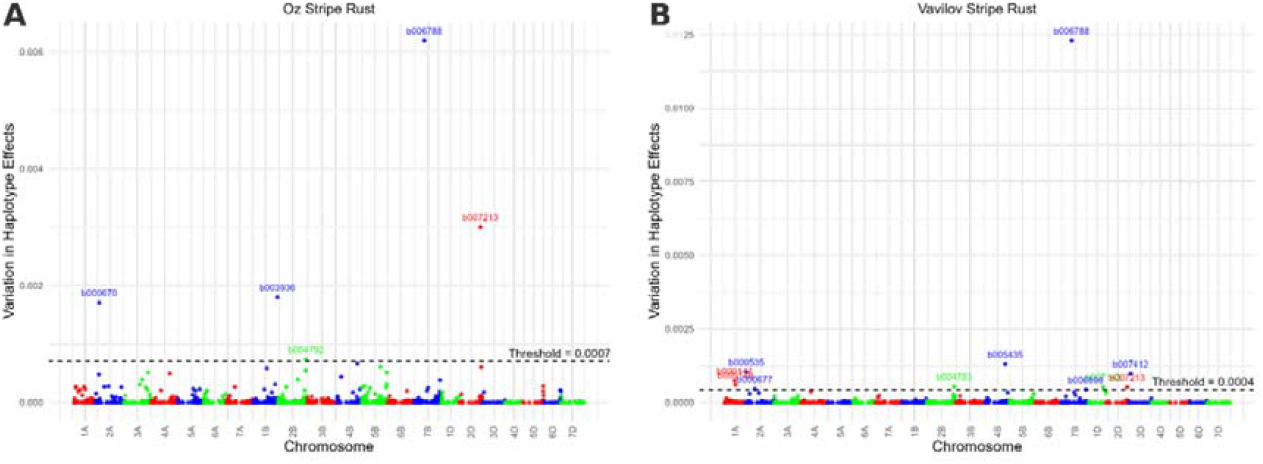
Manhattan plot for stripe rust resistance effects for the top 100 haplotype blocks in the OzWheat (left, A) and Vavilov (right, B) panels. Each point represents a haplotype block; the x-axis is chromosomal position, and the y-axis is the variance of haplotype effects on stripe rust infection. The horizontal dashed line denotes the significance threshold (100 times the genome-wide average variance). Blocks exceeding this threshold are labelled .

Notable regions, 7B:b006788 and 2D:b007213, were highly significant in both populations and represent key loci for stripe resistance (Fig. 5). For leaf rust (using Shenton trial data), under the same stringent threshold, two overlapping haplotype blocks were identified in both the Vavilov and Oz panels: 1A:b000147 and 1B:b003530 (Supplementary Fig. S8a & 8b). Additionally, 7B:b006788 and 2D:b007213 were associated with leaf rust in the Vavilov panel only (Supplementary Fig. S8b); these regions did not reach significance for leaf rust in OzWheat under the same criteria. The co-localisation of these leaf-rust signals with stripe-rust loci suggests either pleiotropy or closely linked resistance genes.

A haplotype resistance catalogue was developed for the OzWheat panel, which includes detailed information on genome-wide haplotype blocks and their corresponding haplotype effect variances for stripe and leaf rust resistance (Supplementary Data). Using this, we also identified five genomic regions with high variances for both diseases across both panels (Fig. 6). We also mapped the top 100 haploblocks for each disease across the three wheat subgenomes (Fig. 6 and Supplementary Fig. S9).

**Fig. 6:**
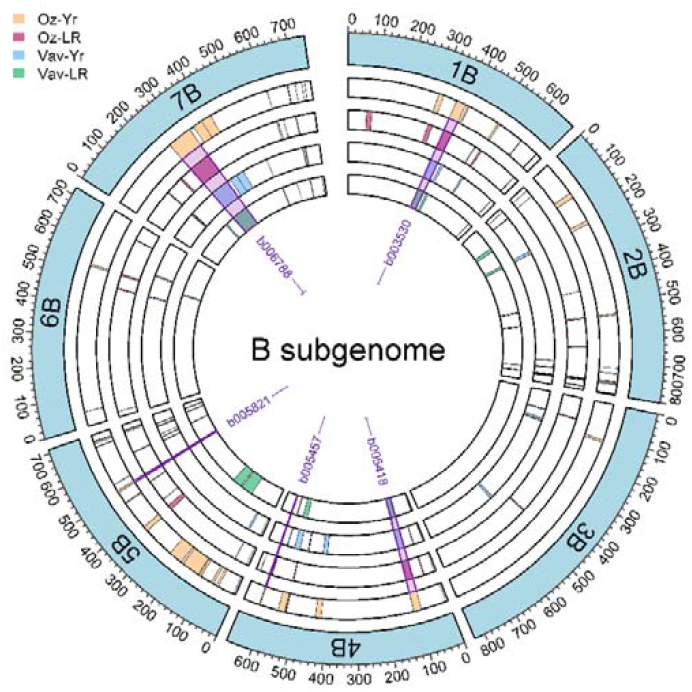
Circos plot showing the chromosome distributions of the top 100 haploblocks associated with stripe and leaf rust resistance. Only B subgenome is displayed here. Five key haploblocks linked to resistance against both diseases are highlighted with purple shading.

Favourable haplotypes were defined as those with negative effects on the rust BLUE (i.e. lower the disease score). The number of favourable haplotypes per line was strongly correlated with resistance. For stripe rust, Pearson’s *r* = -0.76 (95% CI -0.806 to -0.741; *p* = 1.36 × 10 □ ^11^ □), indicating lines carrying more favourable haplotypes had markedly lower infection scores (Fig. 7). A simple classifier using only the favourable haplotypes count achieved AUC =0.87 for discriminating resistant vs susceptible lines. It indicates that there is an 87% chance that a randomly selected resistant individual has a more favourable haplotype profile than a susceptible one (Fig. 7). Concordance of effect estimates between panels differed by trait. For stripe rust, shared haplotypes showed moderate agreement (*r* = 0.55) between OzWheat and the Vavilov core set, suggesting partially conserved genetic architecture. For leaf rust, concordance was low (r = 0.04), pointing to panel-specific haplotype effects despite broadly similar LD architecture. Differences in allele and haplotype frequencies and the use of single site likely reduced power and increased shrinkage in RR-BLUP estimates. This led to many effects being poorly estimated or attenuated toward zero (Supplementary Fig. S7). This highlights that favourable haplotypes in one gene pool may be neutral or unfavourable in another.

**Fig. 7:**
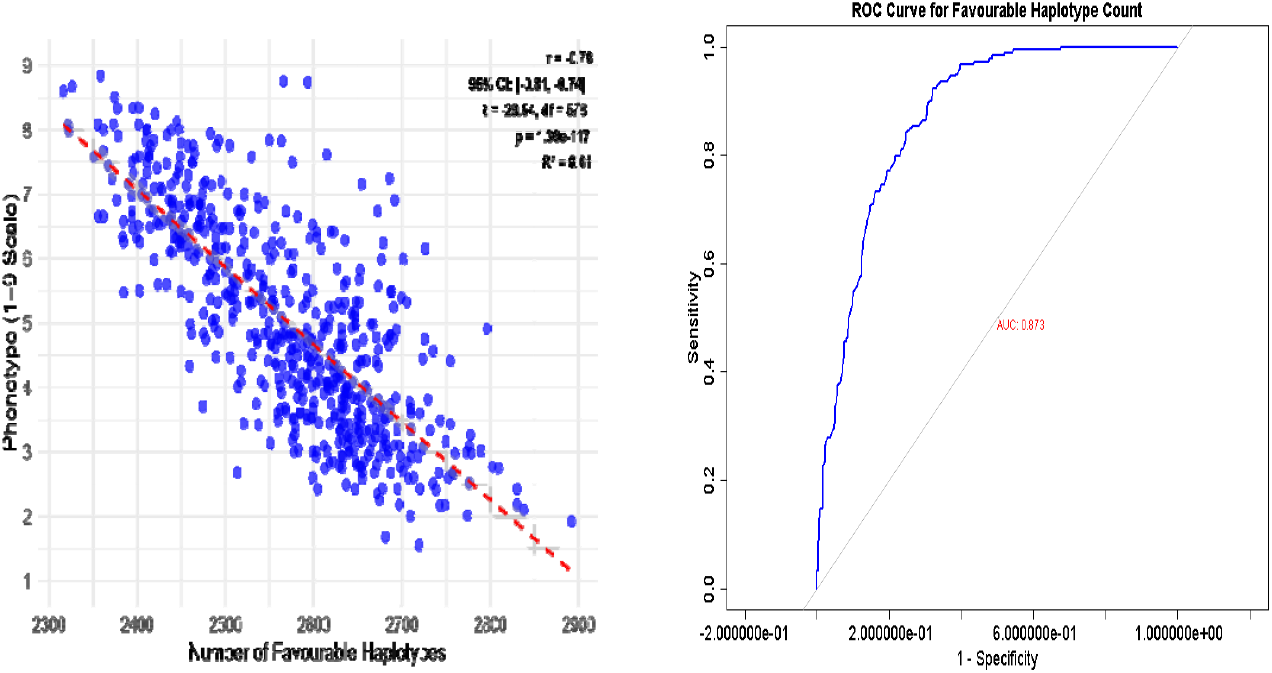
Relationship between favourable haplotype count and phenotypic resistance scores, and Receiver Operating Characteristic (ROC) curve showing the discriminative ability of favourable haplotype count for classifying resistant lines. x-axis (1 – Specificity): represents the false positive rate. As it increases, the classifier becomes more permissive. y-axis (Sensitivity): represents the true positive rate, i.e. the proportion of resistant individuals correctly classified.

#### Haplotype diversity between OzWheat and Vavilov panel

Our unified approach enables direct comparison of haplotype diversity between elite and exotic germplasm. Shannon diversity indices revealed significantly higher haplotype diversity in the Vavilov subset compared to OzWheat (=584), despite its smaller sample size (Wilcoxon rank-sum *W* = 2560.5, *p* < 2.2 × 10^−16^, Supplementary Fig. S10). This confirms landraces as a rich source of novel allelic combinations. Across the combined set of haplotypes, over 19,000 haplotypes were shared between panels, while the Vavilov panel contributed 1,342 unique haplotypes, including 642 favourable haplotypes for stripe rust and 695 for leaf rust that were absent in OzWheat panel. These represent valuable candidates for introgression into modern cultivars.

#### Genetic architecture of stripe rust resistance

To investigate the genetic architecture of stripe rust resistance, genetic variance partitioning was performed using the multi-allelic haplotype model. In this approach, each haplotype block was treated as a locus and haplotypes considered as alleles. The model estimates the proportion of total genetic variance explained by all haplotypes fitted simultaneously. The analysis was carried out using the stripe rust dataset, which exhibited the most reliable phenotypic consistency across trials. Haplotypes alone captured approximately 80% of the total genetic variance, outperforming the SNP-only model (Fig. 8). The combined model did not increase the variance explained much beyond the haplotype model alone, indicating that most of the relevant genetic signal was already captured by haplotypes, and the SNPs only provided redundant information (Fig. 8).

**Fig. 8:**
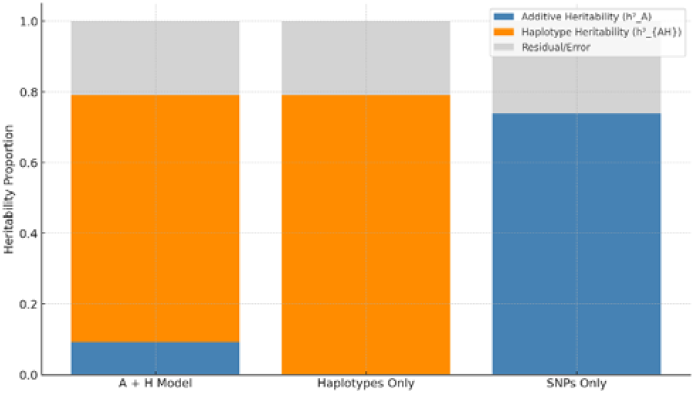
Proportion of heritability explained by different models: SNPs only, haplotypes only, and combined SNP + haplotype model. Haplotypes consistently explain a larger fraction of the total heritability.

To identify genomic regions with large effects, we examined the variance explained by individual haplotype blocks. Each haploblock’s contribution (in standard deviation units) across the genome is visualised in Supplementary Fig. (S11). A few blocks on chromosome 2B and 7B showed relatively higher contributions; however, none exceeded ∼5% of the total genetic variance. This pattern likely reflects the polygenic nature of stripe rust resistance, which is governed by numerous small-effect loci rather than a few major QTLs.

### Parental selection using genome-wide haplotype fitness

To improve rust resistance in the elite cultivar Scepter, we identified diverse parental lines carrying favourable haplotypes absent from its genome. Since no single haplotype block explained a substantial proportion of the genetic variance, we adopted a genome-wide selection strategy to introgress multiple haplotypes simultaneously, rather than targeting a few major regions.

Using the novel fitness function, we selected a group of potential parental lines designed to complement Scepter’s genetic background. The selection index balanced stripe and leaf rust equally, and through a genetic algorithm, we identified 50 lines carrying the most favourable haplotype combinations absent in Sceptre. The selected parents (46 from OzWheat and 6 from the Vavilov collections) were dispersed across the diversity space (Fig. 9A), indicating that they represent a broad genetic background and are not clustered within a narrow subgroup. To further validate their potential, we assessed the GEBVs of these lines for both stripe and leaf rust resistance (Fig. 9B).

**Fig. 9.**
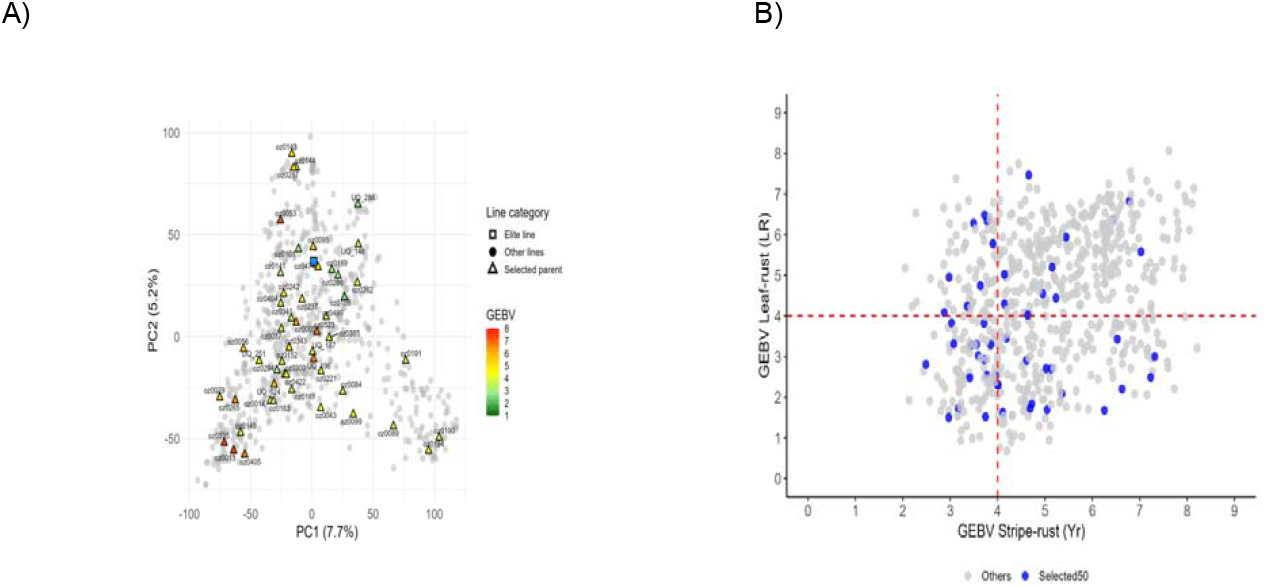
A): PCA plot showing the genetic diversity of the combined panel (OzWheat + Vavilov core set) based on SNP markers. Selected parents (triangles), elite lines (squares, in blue), and other lines (circles) are colored by their GEBV for composite (stripe + leaf) rust resistance. The selected lines span a wide diversity space, reflecting the genetic breadth of the chosen parents. B): Scatterplot of GEBVs for stripe rust (x-axis) and leaf rust (y-axis). Red dashed lines mark the resistance score (more than four tends to be susceptible) for both traits. Selected parental lines (blue) are not limited to the low-left quadrant, illustrating that GEBV alone was not the sole selection criterion; some lines with lower GEBV (susceptible or moderately susceptible) were retained for their resistance haplotypes.

Importantly, while some selected lines ranked high in breeding value (lower is better for resistance) for both traits, they also included some lines classified as susceptible. This result underscores the strategic selection focus on resistance-associated haplotypes, rather than on predicted resistance performance alone. In other words, some lines with poor breeding values were retained because they carry favourable haplotypes which are absent in Scepter, making them valuable donor lines in breeding.

To visualise how the selected parents complement Scepter, a haplotype-state heatmap across resistance-associated blocks was plotted (Fig. 10). The pattern is highly mosaic: no single donor carries the Scepter-alternative state at all blocks, and coverage of candidate states differs among donors. Several blocks are represented by multiple donors, whereas others appear rare (observed in one or two lines). These observations indicate that multiple, complementary parents would be required to capture the full set of candidate regions used in *RustHapSelect*.

**Fig. 10:**
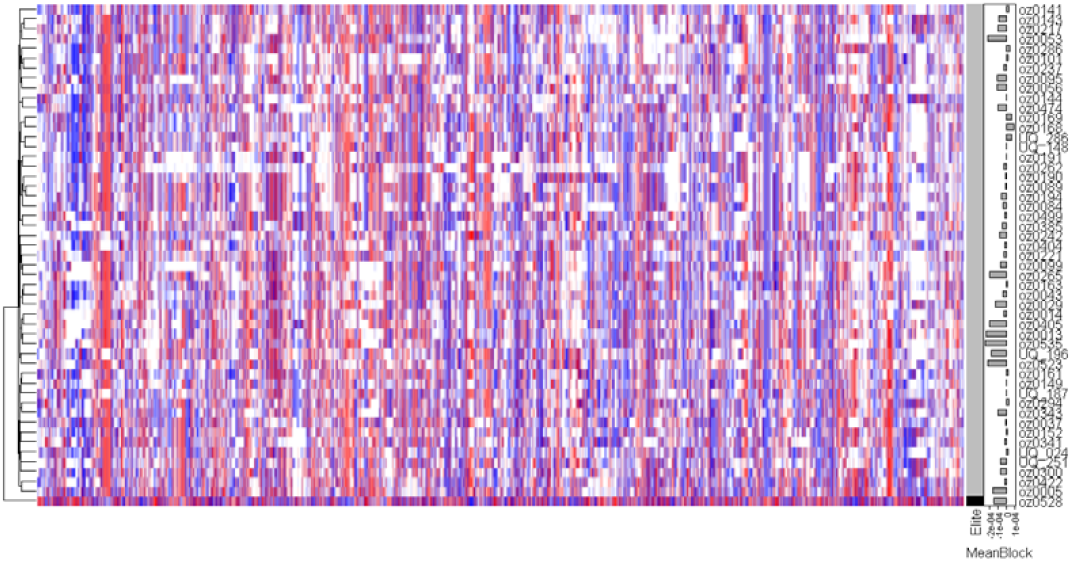
Haplotype states across resistance associated blocks in selected lines relative to ‘Scepter’. Rows correspond to donor or elite lines and columns to candidate haplotype blocks (ordered by genomic position). Blue indicates favourable haplotype and red indicates unfavourable. The heatmap shows relative haplotype effects, using Scepter (or Oz0528) as the baseline genotype for comparison. The righthand bars indicate the mean haplotype effects per line; lower value corresponds to greater resistance.

#### Simulation of progeny performance under breeding strategies

The distribution of mean genetic values (stripe and leaf scores) across breeding generations is shown in Fig. 10 under two contrasting breeding scenarios. In the first scenario, DC progenies were directly advanced through three generations of selfing. As expected, the F_1_ generation exhibited intermediate values with broad variation, reflecting the segregation of alleles introduced from donor lines. The DC generation showed a rightward shift in the distribution, suggesting cumulative gains from combining favourable haplotypes derived from two donor sources. Notably, the Self3 (corresponding to F_4_) generation displayed a narrower, skewed distribution toward higher genetic value, indicating successful fixation of beneficial alleles through selection during selfing (Fig. 11). In contrast, the second scenario involved an additional backcross of the DC lines to the recurrent parent (Scepter) before selfing. The resulting BCSelf3 distribution remained centred closer to zero with reduced variance, suggesting partial recovery of the recurrent parent genome but also a potential dilution of donor-derived resistance alleles (Fig. 11). This comparison highlights a key trade-off: while direct selfing of DCs enables greater genetic gain for rust resistance, incorporating a backcross step enhances background recovery at the expense of genetic diversity and selection potential.

**Fig. 11:**
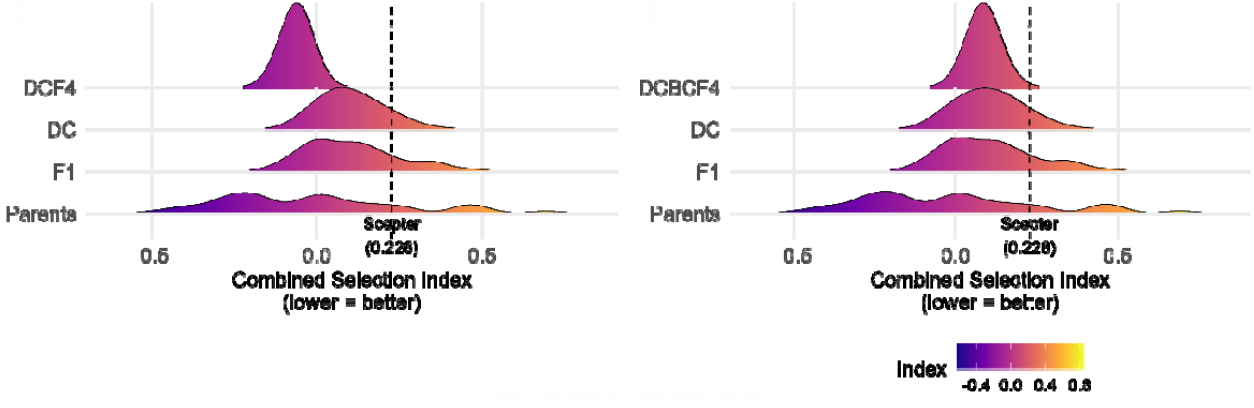
Density distribution of total genetic values for rust resistance across simulated breeding generations under two breeding strategies. The left panel illustrates Scenario 1, where DC progenies were directly selfed for three generations. The right panel represents Scenario 2, where DC progenies were first backcrossed to the recurrent parent Scepter and then selfed for three generations. In both scenarios, each successive generation shows a shift and narrowing of the distribution, with the final selfed generation enriched for resistance-associated genetic values. Colour gradient reflects the density of individuals across the genetic value spectrum. Vertical dashed line indicates the Scepter combined resistance index value for leaf and stripe rust.

## Discussion

This study presents a comprehensive haplotype-based dissection of stripe and leaf rust resistance in wheat, leveraging both elite breeding lines and exotic landraces within a unified framework. By capitalising on high-density genotypic data, multi-environmental phenotyping and genomic prediction, we outline a robust and scalable strategy for identifying and deploying resistance haplotypes in breeding. Focusing on the elite cultivar ‘Scepter’, we show how haplotype-informed selection enhances rust resistance while retaining elite genomic backgrounds. Our results highlight the polygenic architecture of APR and demonstrate how landrace diversity can be effectively harnessed for modern wheat improvement.

### High-quality phenotyping underpins genetic resolution

Robust phenotyping across diverse Australian environments enabled the identification of reproducible genetic signals for stripe rust resistance. High broad-sense heritability (H^2^ > 0.85) across trials confirmed the reliability of stripe rust data, crucial for accurate genetic analysis. In contrast, leaf rust phenotyping was more variable, due to stripe rust co-infection, low disease pressure at some sites, and environmental noise. The Shenton Park (WA-DPIRD) trial, conducted under controlled conditions with known leaf rust pathotypes and with no background natural disease pressure from stripe rust, provided a “clean” dataset with high heritability, proving critical for downstream analysis. This underscores a fundamental principle in breeding for durable resistance: genotypes must be evaluated under conditions that consistently express the target trait. Without targeted screening, key genetic effects may remain undetected due to environmental masking or inconsistent disease pressure (Poland and Rutkoski 2016).

Our observation that location effects outweighed year effects suggests local pathogen populations and microclimates strongly influence rust severity. For example, in 2024, the genotype correlation between sites dropped, particularly involving the Cobbitty trial site. Although pathotype data for this trial were not available, dominant stripe rust strains differed across sites: for instance, pathotype 239 E237 A-17+33+ was present in Victoria (AgVic trials), while 238 E191 A+17+33+ was identified in Queensland. This supports the interpretation that local pathotype variation contributed to differential APR expression. Certain lines may harbour race-specific resistance effective only under particular pathogen pressures, or they may experience delayed resistance onset in cooler microclimates.

To target resistance effective across environments, we estimated haplotype effects using across-environment BLUEs, approximating main effects. This approach highlights haplotypes with consistent effects across sites while reducing site-specific noise. However, it may mask environment-specific responses, especially for traits like disease resistance that are sensitive to local pathogen populations and microclimates. Relying on a single site or year can also be misleading when pathogen populations or environmental conditions differ widely. Future work could explore environment-specific haplotype effects to better capture G×E interactions and race-specific responses. The high-quality of stripe rust datasets gave us confidence in the mapping results and illustrated the value of coordinated multi-institution efforts to generate robust field data. This serves as a reminder that big genetic gains depend on large-scale, well-designed phenotyping (Cobb et al. 2019).

### Haplotype-based mapping enhances resolution and QTL discovery

The haplotype-based association approach used in this study offers improved statistical power over single-marker GWAS. By modelling SNPs within haplotype blocks as joint predictors, it better captures local linkage and reduces the risk of spurious associations from long-range LD. Unlike single-SNP tests, which often suffer from low allelic diversity (Da 2015) and noisy estimates (Tong et al. 2024), haplotypes leverage the collective signal of multiple markers. Furthermore, use of RR-BLUP, which fits all markers simultaneously, accounts for genome-wide LD structure and minimises false positives by distributing effect sizes across correlated loci. This shrinkage-based framework prevents overestimation in high-LD regions, improves the reliability of detected associations.

We identified 7,659 haplotype blocks across the wheat genome, serving as multi-allelic loci in association analysis. Summing SNP effects within blocks to compute local GEBVs proved effective for pinpointing trait-associated loci. This aligns with growing evidence that haplotypes better capture functional inheritance in polyploid crops like wheat (Bhat et al. 2021; Brinton et al. 2020; Voss-Fels et al. 2019). By examining haplotype-effect variance within blocks, we flagged loci with different allele effects, indicative of a segregating QTL. This variance-based approach, which tests whether any haplotype within a block differs largely in effect, reduces the burden of multiple testing and is less sensitive to MAF issues, as rare variants are aggregated into common haplotypes (Kemper et al. 2015; Xiang et al. 2021).

Our results showed good concordance with known rust resistance loci. For stripe rust, haplotype block 1BL:b003936 (∼ 669.4 Mb) corresponds to the well-characterised *Lr46*/*Yr29* region, a source of partial, non-race-specific APR to multiple rusts and powdery mildew. This region, originally identified in cultivar Pavon 76, is also associated with the *Ltn2* leaf tip necrosis trait (Li et al. 2024), and we detected the corresponding haplotype in both the OzWheat and Vavilov panels. Another notable block, 7B:b006788 (285.7 Mb), appeared in both panels for stripe rust (and in Vavilov for leaf rust). It lies within the reported interval of *QYr*.*caas-7B* (145-337 Mb), an APR QTL for stripe rust that is independent of the known *Yr041133* gene on 7BL (Li et al. 2024). Our block lies near the centre of this interval, suggesting it represents the same or a closely linked locus. The distal region of 7BL (∼ 608-722 Mb) is a known hotspot for rust resistance that includes genes *Lr14a* (Ren et al. 2023) and *Lr68* (Herrera-Foessel et al. 2012; Wang et al. 2025). From the Shenton Park leaf rust trial, block 7BL:b006898 overlapped with *Lr14a*, a race-specific gene encoding an ankyrin repeat protein, though this signal was evident only in OzWheat. Additionally, 7B:b006880 (632.6 Mb) aligned with the *Lr68* region, a “slow-rusting” APR gene originally mapped using SSR markers. Block 7B:b007020 (721.2 Mb) block detected in both populations for stripe rust also lies in this region of 7BL. Although no major stripe rust gene has been definitively mapped to this region, a recent study by Wang et al. (2025) identified a ∼30Mb segment on 7BL (715.77-733.25 Mb) in Chinese cultivar Aikang58 harbouring multiple loci (*YrAK58*.*1, YrAK58*.*2*, and *YrAK58*.*3*) that contribute to durable resistance against stripe rust. The positional overlap and similar APR expression patterns suggest that 7B:b007020 may capture signals from one or more of the *YrAK58* loci. Importantly, the Vavilov-specific block, 7B:b006898, aligns with the known *QYr7B*.*2* (*Yr39*) locus (Zheng et al. 2022), a gene conferring durable high-temperature APR to stripe rust. These results suggest that the long arm of 7B harbours multiple APR loci contributing to both stripe and leaf rust resistance, with evidence of pleiotropy and environment-specific expression. Detection across independent germplasm sets reinforces the biological relevance and robustness of our associations.

In addition to known loci, we also identified regions with no prior documentation, indicating potential novel resistance sources. Notably, 2D:b007213 showed strong association with stripe rust in both panels, yet no known *Yr* gene or QTL has been reported at this position. We also identified panel-specific regions, such as 2A:b000535, in the Vavilov panel, which overlaps multiple previously reported stripe rust QTLs, *QYr*.*uga-2AS_26R61* (Hao et al. 2011), *QYr*.*ucw-2AS_PI610750* (Lowe et al. 2011), *QYr*.*wpg-2A*.*2_IWA8274* (Ward et al. 2021), supporting its relevance. These novel or less-documented resistance loci provide new opportunities to broaden the genetic base for rust resistance.

### Haplotype catalogue and validation across elite and exotic germplasm

This study demonstrates how haplotype-based mapping in elite wheat lines, combined with validation using exotic landraces, can inform breeding decisions for durable rust resistance. Primary association mapping was conducted in the OzWheat panel, where high-resolution haplotypes linked to stripe and leaf rust resistance were identified and catalogued. Validation with 50 Vavilov landrace lines revealed 1,342 unique haplotypes (642 favourable for stripe rust, 695 for leaf rust) that were not present in OzWheat, highlighting the value of historical germplasm for uncovering novel resistance.

Importantly, certain landraces emerged as especially valuable based on their genetic diversity and resistance potential. For example, accessions UQ_024 and UQ_187 ranked highest for haplotype diversity (Shannon index), indicating they carry a broad array of haplotypes. These same lines were also prioritised by the introgression fitness index, which balanced stripe and leaf rust breeding values, reinforcing their value as promising donor ideotypes: not necessarily agronomically optimal themselves, but they are enriched in resistance haplotypes that are underrepresented in elite breeding pools. Most favourable haplotypes had small effects, consistent with a polygenic resistance model. This supports a shift toward haplotype stacking and genomic selection, as conventional phenotypic selection struggles to accumulate minor-effect alleles efficiently. Notably, among the 52 stripe rust-associated regions, 15 were exclusive to landraces, with several overlapping (2B: b004792, 6A: b002416) known QTLs (Jambuthenne et al. 2022; Riaz et al. 2018; Tong et al. 2024). This underscores that landraces harbour many valuable resistance loci that have been lost or never introduced into modern wheat. We also identified haploblocks conferring resistance to both stripe and leaf rust (e.g. 7B:b006788 and 2D: b007213), suggesting pleiotropy or shared defence mechanisms. Candidate genes in these regions may involve signalling, structural defence, or hormone-regulation pathways as described by Hafeez et al. (2021) and Song et al. (2023), offering targets for future functional validation.

The polygenic nature of APR and the genomic complexity of polyploid wheat pose challenges for deploying individual resistance genes. Haplotype-based analysis provides a practical alternative by capturing co-inherited blocks of favourable alleles that function as meaningful selection units in breeding. In this study, we identified high-value haploblocks, while marker development was beyond the scope of this work, these regions define genomic intervals suitable for future development of KASP assay targeting high-effect haplotypes to enable marker-assisted selection. Importantly, targeting haploblocks with multiple minor-effect loci may be more effective than pyramiding isolated resistance genes, especially for complex traits like APR. Examples like the *Rht-B1b–EamA-B–ZnF-B* haploblock (Song et al. 2023) contribute to yield, nitrogen use efficiency, and plant architecture, demonstrating the pleiotropic potential. Conversely, combining *Yr29* with other APR loci in cultivars like Jing 411 (Li et al. 2024) exemplifies haplotype stacking. Together, these examples highlight complementary strategies: leveraging pleiotropic haploblocks and stacking favourable haplotypes to enhance trait performance and durability of resistance.

### Creating pre-breeding pools enriched for rust resistance haplotypes

To develop pre-breeding pools enriched for rust resistance while retaining elite genomic backgrounds, we prioritised favourable haplotypes absent from the benchmark cultivar ‘Scepter’. This fitness function mirrors genomic selection indices but is specifically tailored to capture complementary alleles relative to a given elite line. In essence, the selection strategy focuses on favourable haplotypes over absolute resistance levels, meaning that even a line with moderate susceptibility can still be valuable if it carries unique resistance haplotypes. Simulations using AlphaSimR showed that stacking complementary haplotypes through targeted crosses using forward breeding or backcrossing can increase resistance while preserving elite backgrounds to some degree. These findings align with recent advances advocating haplotype-informed crossing for polygenic trait improvement (Tong et al. 2024; Voss-Fels et al. 2019).

### Limitations and Future Directions

While this study establishes a haplotype-informed foundation for breeding against stripe and leaf rust, several limitations remain. First, we phenotyped the 50-line Vavilov core rather than the full collection. This was a resource-driven design choice, which reduces statistical power and may underrepresent rare haplotypes. Expanding to the full Vavilov collection will increase resolution and uncover additional resistance loci, particularly rare but potentially valuable ones. Second, the use of an additive RR-BLUP model does not capture non-additive effects or genotype-by-environment (G×E) interactions. Future work could incorporate non-linear or epistatic interactions into models to better reflect the complexity of APR genetics. Additionally, the resolution of candidate regions could be improved using higher-density marker platforms or long-read sequencing to refine LD blocks and narrow down causal variants. A further challenge is characterising genotype × environment × pathogen (G×E×P) interactions. Although our dataset spans multiple agro-ecological zones and local rust pathotypes, deeper pathotype-specific screening and integration of high-throughput phenotyping will improve trait dissection. Combining transcriptomics or expression QTL (eQTL) data may also help prioritise functional variants within haploblocks, accelerating the path from association to application.

### Conclusion

By identifying and quantifying key rust resistance haplotypes and applying them in a predictive breeding framework, this work provides both conceptual and practical tools for developing rust-resilient wheat cultivars. The methodology is broadly applicable and can be extended to other complex traits and crop–pathogen systems, contributing to more informed and sustainable crop improvement strategies. Overall, this study provides a unified framework for identifying, validating, and deploying resistance haplotypes across elite and exotic material. By combining mapping, diversity analysis, fitness-based prioritisation, and simulation, we outline a practical strategy for developing polygenic rust resistance in wheat. This streamlined approach exemplifies how pre-breeding and applied breeding can be unified.

## Supplementary data

Fig. S1. PCA plot of the full Vavilov panel and labeling selected core set

Fig. S2. Comparison of allele frequencies between OzWheatV2 and OzWheatV1 (used as reference panel)

Fig. S3. Comparison of allele frequencies between Vavilov and OzWheatV1 (used as reference panel)

Fig. S4. Percent of missing phenotypic data for leaf and stripe rust across location and years

Fig. S5. Broad-sense heritability estimates for leaf and stripe rust across location

Fig. S6. LD decay in OzWheat and Vavilov panel

Fig. S7. Distribution of SNP effects in OzWheat and Vavilov for stripe and leaf rust Fig. S8. Manhattan plot for leaf rust in OzWheat and Vavilov panel

Fig. S9. Chromosomal distribution of the top 100 haploblocks associated with stripe and leaf rust resistance across the A and D subgenomes

Fig. S10. Haplotype diversity between OzWheat and Vavilov core set

Fig. S11. Standardised heritability estimates for each haploblock across the genome

Table S1. Descriptive statistics for leaf and stripe rust severity scores across location and years

Table S2. Distribution of 29,972 markers across chromosomes A, B, and D from different genotyping platforms

## Acknowledgements

The authors acknowledge Melania Figueroa and Jana Sperschneider for their ongoing contributions to wheat rust research and for providing valuable insights that strengthened this work.

## Author contributions

SY: data curation, investigation, data analysis, writing-original draft; SY, BJH, LTH, PND: conceptualisation and study design; SD, MM: genotypic data generation; RM, PND: phenotypic data generation; SY, SD, MM, ED, PND, BJH, DJ, LTH, RM: writing-review and editing; PND, LTH, BJH: funding acquisition.

## Conflict of interest

The authors declare that they have no conflicts of interest in relation to this work.

## Funding

Funding was provided by Grains Research and Development Corporation, CSP2304-013RTX.

## Data availability

All data supported the findings of this study are available through the CSIRO Data Access Portal (DAP, https://data.csiro.au/collection/csiro:61604).

## Notes

### Competing Interest Statement

The authors have declared no competing interest.

